# Single-cell RNA sequencing of CTLA-4 and PD-1 blockade in pulmonary paracoccidioidomycosis highlights a protective transcriptional program mediated by activated Th17 cells, neutrophils and macrophages

**DOI:** 10.64898/2026.01.21.700363

**Authors:** Sandra Marcia Muxel, Dennyson Leandro M. Fonseca, Ian Antunes Ferreira Bahia, Patrick da Silva, Leonardo Mandu-Gonçalves, Emanuella Sarmento, Jonathan Miguel Zanatta, Clara Andrade Teixeira, Kathia Terumi Kato, Manoela Osorio Reis Sales, Carlos Neandro Cordeiro Lima, Bruno Prado Eleuterio, Luciana dos Santos Barros Manhães, Bernardo de Castro Oliveira, Nycolas Willian Preite, Bruno Montanari Borges, Valeria de Lima Kaminski, Marina Cacador Ayupe, Bianca Vieira dos Santos, Maria Regina D’Império Lima, Denise Morais da Fonseca, Flavio Vieira Loures, Vera Lúcia Garcia Calich

**Affiliations:** Laboratory of Molecular Immunology Regulation miRlab, School of Arts, Science and Humanities (EACH, University of São Paulo (USP), São Paulo, SP, Brazil; Postgraduate Program on Immunology, Institute of Biomedical Science (ICB), University of Sao Paulo (USP), Sao Paulo, SP, Brazil; Brigham Women’s Hospital - Harvard Medical School, USA; Laboratory of Mucosal Immunology - Department of Immunology - Institute of Biomedical Sciences, University of Sao Paulo, São Paulo, Brazil; Laboratory of Infectious Diseases Immunology - Department of Immunology - Institute of Biomedical Sciences, University of Sao Paulo, São Paulo, Brazil; Immune Health Laboratory, “Regulation of host responses and immune health” (IRL2029), French National Centre for Scientific Research (CNRS) and Ribeirão Preto Medical School (FMRP) of the University of São Paulo (USP); National Institute of Science and Technology for Research in Infectious and Chronic Diseases of the Mucosa and Skin; Institute of Science and Technology, Federal University of São Paulo, José dos Campos, SP, Brazil; Laboratory of Mycosis Immunology - Department of Immunology - Institute of Biomedical Sciences, University of Sao Paulo, São Paulo, Brazil

## Abstract

Pulmonary paracoccidioidomycosis (PCM) relies on a finely balanced lung-immune network in which Th17, Treg, neutrophils, and macrophages orchestrate fungal control and tissue integrity. Furthermore, within the context of single-cell sequencing, little is known about how immune checkpoint inhibition modulates this balance during systemic mycosis. In a previous study we verified that the blockade of checkpoint molecules (CTLA-4 and PD-1) restores protective immunity that reduces fungal loads, tissue pathology and mortality of infected mice. Here, we have further studied this model by single-cell RNA sequencing on lung leukocytes from mice infected with *Paracoccidioides brasiliensis* and treated with anti-CTLA-4 or anti-PD-1 antibodies to define the cellular and molecular consequences of checkpoint blockade in vivo. We generated a high-resolution atlas covering T cells, neutrophils, macrophages/monocytes, B cells, NK cells, and epithelial subsets. Checkpoint inhibition consistently remodeled the CD4⁺ T cell compartment toward a Th17-enriched program, with reduced interleukin-10 expressing Treg frequencies and a prominent interleukin-17A⁺ (IL-17), C-C chemokine receptor type 2⁺, CXC chemokine (CXC) receptor type 6⁺, CD44⁺ (*Il17a*⁺*Ccr2*⁺*Cxcr6*⁺*Cd44*⁺) signature. This shift was accompanied by the expansion and activation of neutrophils and macrophages expressing tumor necrosis factor and chemokines (including CXC motif chemokine ligand (*Cxcl*1, *Cxcl2* and *Ccl4*), microbicidal-associated genes (S100 calcium-binding protein A8 and A9), and regulatory mediators such as secretory leukocyte protease inhibitor and interleukin-15. Ligand–receptor and trajectory analyses revealed a reinforced Th17–myeloid communication axis, particularly under CTLA-4 blockade, converging on IL17–centered inflammatory circuits while attenuating canonical Treg checkpoints (cytotoxic T-lymphocyte-associated protein 4, programmed cell death protein 1, interleukin-10). Together, these data demonstrate that anti-CTLA-4 and anti-PD-1 immunotherapies profoundly rewire the lung-immune microenvironment during PCM, amplifying effector Th17–neutrophil–macrophage networks that benefits protective antifungal activity Our work provides a mechanistic framework for evaluating immune checkpoint inhibitors in chronic fungal infections.

## INTRODUCTION

Pulmonary paracoccidioidomycosis (PCM) is a subacute systemic mycosis caused by the thermically dimorphic fungus *Paracoccidioides brasiliensis*, *P. lutzii*, *P. americana*, *P. venezuelensis*, and *P. restrepiensis*, affecting an estimated ten million individuals in Latin America ^1–4^. Despite widespread exposure, only 1-2% of infected people develop clinically apparent disease^5,6^. Infection begins with the inhalation of airborne conidia that reach the alveoli and are phagocytosed by resident macrophages; within days, they transition to the pathogenic yeast form and proliferate^7–10^. The pulmonary presentation often resembles tuberculosis or malignancy, and chronic forms are characterized by progressive lung involvement^11,12^. Th17 cells, a major source of IL-17A and IL-22, are central to antifungal defense by promoting antimicrobial peptide production and neutrophil recruitment in the^11–16^. The *P. brasiliensis* can exploit regulatory pathways, and PCM is marked by accumulation of Tregs expressing CTLA-4 and PD-1, alongside elevated levels of the cytokines TGF-β and IL-10, which collectively suppress protective immunity^14–20^. Immune checkpoint inhibitors, targeting PD-1 and CTLA-4, can reverse such inhibitory signaling and have transformed immunotherapy^21–25^. Our previous study has demonstrated that the blockade of these molecules in a pulmonary model of paracoccidioidomycosis exerts a protective effect as revealed by enhanced immunity, decrease fungal loads, tissue pathology and mortality of infected mice^26^. However, their effects during infection, the molecular modulation at the single-cell level in the pulmonary environment remain poorly defined^26–28^.

In this study, we applied the single cell RNA sequencing (scRNA-seq) to profile the pulmonary immune landscape during *P. brasiliensis* infection and following treatment with anti-PD-1 and anti-CTLA-4. This approach enables high resolution mapping of CD4^+^ and CD8^+^ T cells, neutrophils, macrophages and additional immune populations, revealing transcriptional programs associated with activated Th17 and Tregs. In addition, by integrating ligand–receptor inference and trajectory analyses, we identified immune circuits underlying IL-17-driven responses and neutrophil activation^23,29,30^. Thus, we applied a systems biology analysis to characterize how *P. brasiliensis* infection interacts with PD-1 and CTLA-4 blockade to reprogram the protective transcriptional architecture of the pulmonary microenvironment.

## RESULTS

### Mapping of lung cellular composition in the lung of murine *P. brasiliensis* infection during immune checkpoint inhibitors treatment

To investigate the pulmonary immune response against fungal infection to *P. brasiliensis* and the impact of treatment with anti-PD-1 and anti-CTLA-4 treatment, we established a murine model of intratracheal infection, such as described^26^. Three weeks after infection, groups of animals were treated with PBS, normal IgG, anti-PD-1, and anti-CTLA-4, three times per week, until the tissue harvesting point at week 8. The lung tissue was then processed for scRNA-seq using the BD Rhapsody WTA platform (**Fig. 1a**).

**Figure 1.**
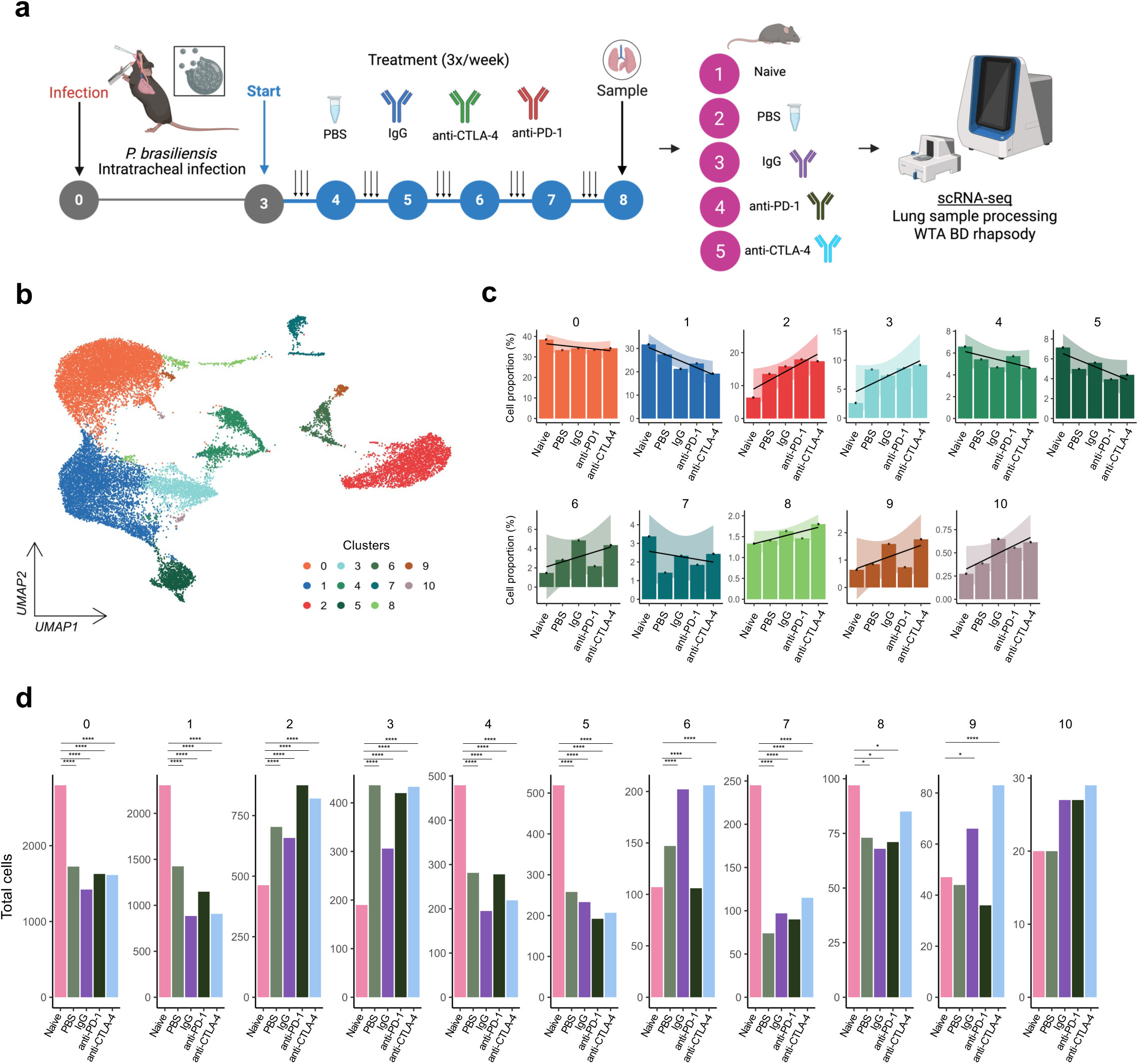
Experimental design and global remodeling of lung leukocyte composition during checkpoint blockade in PCM. **a)** Schematic of the intratracheal infection with *P. brasiliensis* (1×10^6^ Pb18 yeasts) and treatment schedule with PBS, IgG, anti-PD-1, or anti-CTLA-4 groups for lung leukocyte collection and BD Rhapsody WTA scRNA-seq. **b)** UMAP embedding of 70,474 CD45^+^ lung cells colored by the 11 transcriptionally defined clusters. **c)** Barplots show the cell proportions per cluster for each treatment group. The shadows represent the standard error (SE) of the proportion of cells between treatment groups. **d**) Barplots show total of cells with statistical significance between treatment groups. The statistical significance was determined using two-sided Wilcoxon rank-sum tests and is indicated by asterisks (\**p* ≤ D0.05, \*\**p* ≤ 0.01, \*\*\**p* ≤ 0.001, and \*\*\*\**p* ≤ 0.0001).

Using tools available in the Seurat R package^31^, 11 distinct clusters (**Fig. 1b**) with an average of 660, 471, 378, 442 and 429 cells for the Naive, PBS, IgG, anti-PD-1, and anti-CTLA-4 groups, respectively. (**Supp. Table 1**). In this context, we investigated the trend in cell distribution among the experimental groups, quantifying the relative abundance of each cluster among five groups. The analysis revealed dynamic changes in the proportions of the groups in response to treatment (**Fig. 1c** and **Supp. Table 1**). Notably, clusters 0, 1, 4, 5 and 7 showed a downward trend in the treated groups compared to the Naive. In contrast, clusters 2, 3, 6, 8, 9 and 10 showed an upward trend in the same way (**Supp. Fig. 1**). This was corroborated by an analysis of multiple comparisons showing significant variation (adjusted *p* < 0.05) in the number of cells in each cluster in the different groups (**Fig. 1d** and **Supp. Table 2**).

To characterize the cellular signatures present in the lung during infection and treatment, we performed differential expression (**Supp. Table 3**) and biological process (BP) enrichment (**Supp. Table 4**) analyses across all clusters. We identified a distinct cellular characterized by high expression of genes such as *Cd79a*, *Cd19*, *Cd3e*, *S100a8*, *S100a9*, *Hba-a1*, *Nkg7*, *Il1b*, *Cd36*, *Lgals3*, *Tspan7* and others (**Fig. 2a** and **Supp. Fig. 2**). In addition, the set of genes identified together with the markers were enriched and indicated different profiles for each cluster (**Fig. 2b**). Based on these results, we annotated the following cell types: B cells (cluster 0), T cells (clusters 1, 3, and 10), neutrophils (cluster 2), red blood cell precursors (cluster 4), natural killer (NK) cells (cluster 5), macrophages (cluster 6), monocytes (cluster 7), alveolar type II cells (cluster 8), and eosinophils (cluster 9) (**Fig. 2c** and **Supp. Table 5**). Notably, we identified a cluster with a red blood cell profile, which was excluded from subsequent analyses.

**Figure 2.**
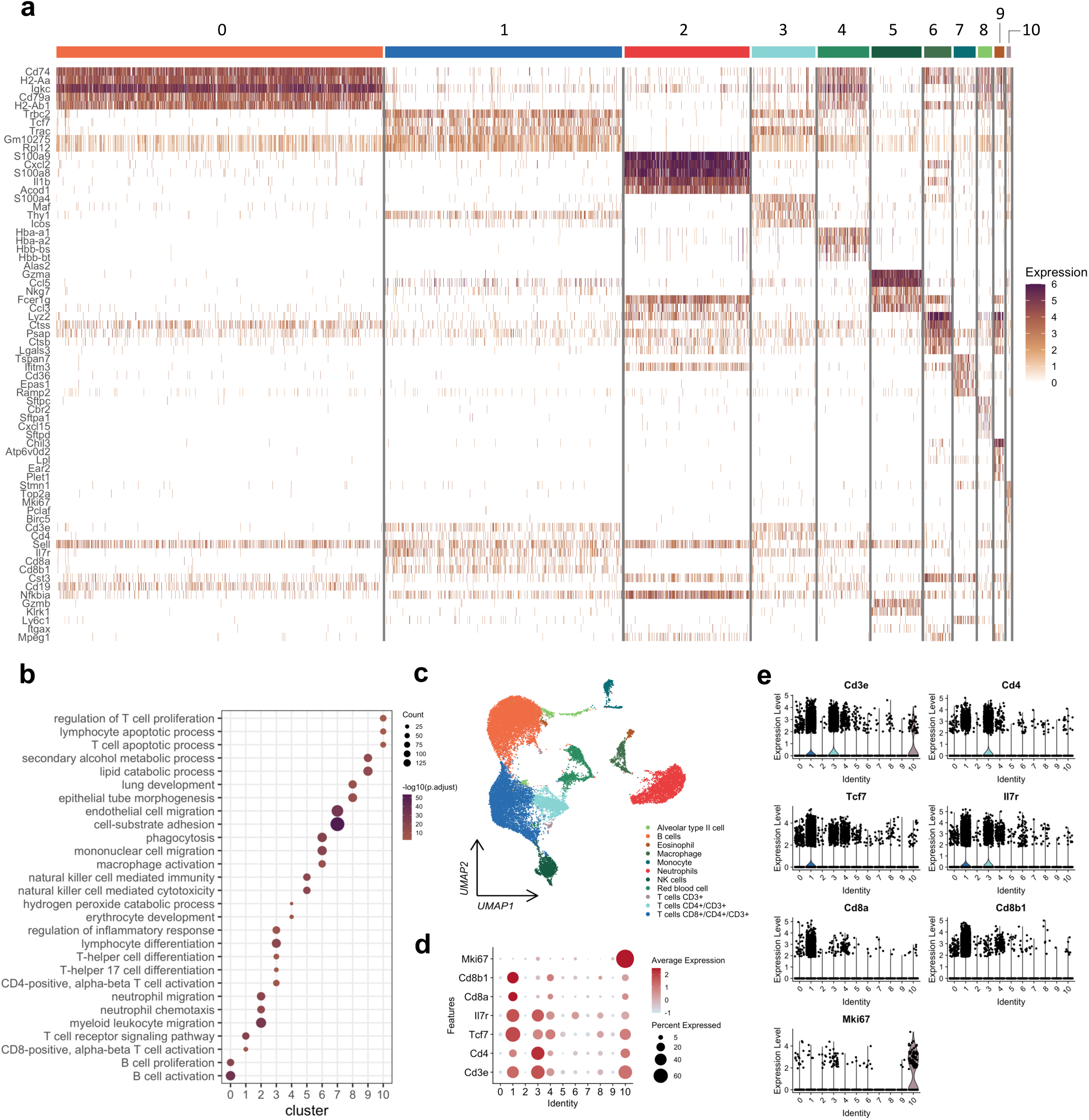
Transcriptional signatures and functional annotation of lung cell populations. **a**) Heatmap of differentially expressed genes in the 11 clusters, highlighting differentially expressed marker genes with lower FDR values. The bars above the heatmap represent the colors of each cluster. **b**) Dotplot showing enriched biological processes based on differentially expressed genes in each cluster. The size of the dot represents the gene count within the pathway, and the color range is represented by the -log10 of FDR. **c**) UMAP represented by annotated cell types. **d-e**) Dot and Violin plots of representative genes per cluster showing expression intensity and fraction of positive cells, confirming the identity of major lineages and activation states.

In more detail, the high expression of T cell–associated genes set, including *Cd3e*, *Cd4*, *Cd8a*, *Cd8b1*, *Tcf7*, *Il7r*, and *Mki67*, was enriched in biological process related to T cell activation, proliferation, and differentiation (**Fig. 2d–e**), indicating the presence of distinct subsets of activated and proliferating T cells. These findings pave the way for the idea that blocking the checkpoint, i.e., anti-CTLA-4 and anti-PD-1, in a *P. brasiliensis*-infected model leads to a marked change in lung immunity, specifically, with greater activation and proliferation of effector CD4^+^ T cells, ultimately improving fungal elimination^26^.

### T cell profile in the lungs of *P. brasiliensis*-infected mice treated with anti-PD-1 and anti-CTLA-4

To further characterize T cells profile in the context of infection and immunotherapy, we clustered only *Cd3e*-positive populations, resulting in five distinct T cell subpopulations (**Supp. Fig. 3a**). Classical T cell markers, including *Cd3e*, *Cd3d*, *Cd4*, *Cd8a*, *Cd8b1*, *Il7r*, *Il2ra*, *Il2rb*, *Thy1*, *Eomes*, *Ifng*, and *Xcl1* were used to define their phenotypes (**Supp. Fig. 3b–c**). Clusters 0 and 1 expressed *Cd8a* and *Cd8b1*, indicating a CD8⁺ T cell profile. Notably, cluster 1 also expressed *Il2rb*, *Eomes*, *Ifng*, and *Xcl1*, suggesting an activated effector phenotype characteristic with cytotoxic CD8⁺ T cells responding to inflammation. Interestingly, cluster 0 co-expressed *Cd4* and *Cd8a*, possibly reflecting a heterogeneous population composed of both CD4⁺ and CD8⁺ T cells. In addition, the clusters 2, 3, and 4 were enriched for *Cd4*, *Il7r*, and *Thy1*, consistent with a CD4⁺ T cell phenotype, with clusters 3 and 4 showing almost exclusive expression of *Cd4*. Importantly, clusters 3 and 4 also showed an increased proportion of cells in the treatment groups (**Supp**. **Fig. 3d** and **Supp. Table 6**), suggesting that immunotherapy promoted the expansion of CD4⁺ T cells. Together, these findings indicate that pulmonary infection followed by immune checkpoint blockade results to increased frequencies of T cells with likely effector and helper phenotypes (**Supp**. **Fig. 3d**), reinforcing a shift toward T cell–mediated immunity under therapeutic conditions.

In this context, we filtered only the clusters 3 and 4 to better characterize the T cell profile. This approach allows us to investigate the dynamics and plasticity of the CD4⁺ T cells in the pulmonary microenvironment. First, we found three new main subpopulations (**Fig. 3a**) which, through a differential expression analysis between the clusters, we identified characteristic markers. Based on the false discovery rate (FDR) values, we observed that cluster 2 obtained a total of 343 differentially expressed genes (DEG). Interestingly, clusters 0 and 1 had a lower number of DEGs, 4 and 45, respectively (**Fig. 3b**).

**Figure 3.**
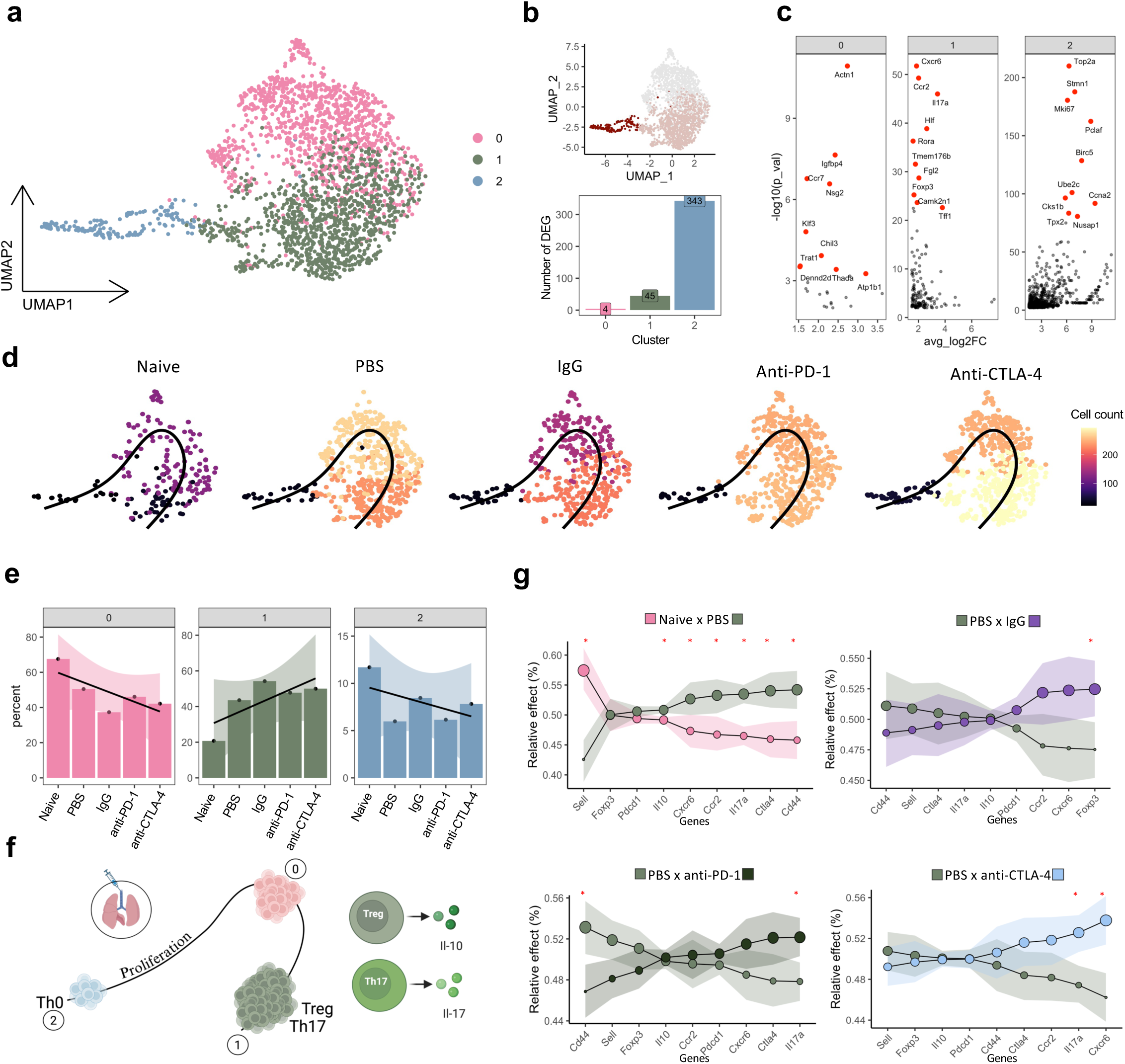
CD4^+^ T cell remodeling with Treg/Th17 plasticity and *Il17a*-biased trajectories under checkpoint blockade. **a)** UMAP of reclustered CD4^+^ T cells revealing three T cell subpopulations. b) UMAP and bar plots representing the number of DEGs for each subcluster, highlighting the highly proliferative set 2 and the mixed Treg/Th17 set 1. **c**) Dotplots illustrating the genes that define each CD4^+^ subgroup. The red dots represent the top markers for each cluster. **d**) UMAP showing the pseudotemporal trajectory, using cluster 2 as the origin and showing the differentiation of proliferating CD4^+^ T cells into Treg/Th17 states throughout infection and treatment. The color spectrum represents the cell count within each cluster. **e**) Barplots show the cell proportions per T cell subcluster for each treatment group. The shadows represent the standard error of the proportion of cells between treatment groups. **f**) Diagram representing the identified trajectory with the Treg and Th17 proliferation phenotype observation, highlighting IL-10 and IL-17. **g**) Relative effect curves for *Il17a* and other key genes under all conditions (Naïve vs PBS, PBS vs IgG, PBS vs anti-PD-1, PBS vs anti-CTLA-4). The shadows represent the 95% confidence interval (CI), and the size of the dots represents the relative increase.

Furthermore, to exemplify gene sets with high expression we highlight DEGs based on *p*-value. This representation allows a clearer visualization of the most significantly modulated genes within each cluster, particularly in clusters 1 and 2. Cluster 1 presents mixed characteristics between regulatory T cells (Treg) and T helper 17 cells (Th17), with expression of *Foxp3*, *Il10*, *Il17a*, *Rora*, *Ccr2*, *Cxc6* and *Hif1a*, suggesting phenotypic plasticity between Treg and Th17. Cluster 2 strongly expresses proliferation-related genes, such as *Mki67*, *Top2a*, *Pcna* and *Stmn1*, indicating a population in a highly proliferative state. Cluster 0, on the other hand, exhibits genes associated with more basal signaling and low inflammatory activity, expressing few genes such as *Actn1*, *Ccr7*, *Igtpbd4* and *Neg2*, suggesting a transition between profiles (**Fig. 3c; Supp. Fig. 4a-c** and **Supp. Table 7).**

Based on these findings and to investigate CD4⁺ T cell differentiation dynamics within the lung microenvironment, we performed trajectory analysis, setting cluster 2 as the starting point and cluster 1 as the endpoint. This analysis revealed that, as infection progresses and treatment is administered, proliferative CD4⁺ T cells (cluster 2) differentiate into Treg and Th17 phenotypes (cluster 1) (**Fig. 3d**). Subsequently, we quantified the distribution of CD4⁺ T cell profiles across the experimental conditions. Notably, cluster 1 exhibited the highest frequencies of cells in the anti-CTLA4 (294 cells) and anti-PD1 (256 cells) groups compared to the control groups: IgG (218), PBS (233), and Naive (39) (**Supp. Table 8**). The cluster distribution further highlighted a relevant shift toward cluster 1 under immune checkpoint blockade (**Fig. 3e** and **Fig. 3f**). Furthermore, we observed a significant increase in *Il17a* expression in both comparisons: PBS vs anti-PD-1 and PBS vs anti-CTLA-4 (**Fig. 3g** and **Supp. Fig. 4d-e**). In both cases, *Il17a* expression was significantly regulated after treatment, highlighting its potential role in modulating the immune response induced by PD-1 and CTLA-4 inhibition^26^.

### Anti-CTLA-4 drives transcriptional cell–cell communication toward an *Il17a* axis in the pulmonary microenvironment upon *P. brasiliensis* infection

To investigate how infection and treatment with immune checkpoint inhibitors (anti-PD-1 and anti-CTLA-4) reorganize cell-cell communication in the pulmonary microenvironment, we inferred ligand–receptor (L-R) interactions between cell types. Initially, communication analysis was conducted on the total set of CD4⁺ T cells. The groups exhibited similar communication pattern between CD4⁺ T cell, macrophage, neutrophil, and B cell populations, which were most evident in PBS, IgG, and anti-CTLA-4 groups (**Fig. 4a**). To test whether the observed communication pattern could be driven by specific subpopulations, we examined Pearson correlations between the frequency of Treg/Th17 and other populations (**Fig. 4b** and **Supp. Table 9**). The results indicated a strong positive correlation of Treg/Th17 with neutrophils and a similar, although more discreet, trend with macrophages. In addition, we observed that this correlation may be influenced mainly by the increase in these populations in the PBS, IgG, anti-PD-1, and anti-CTLA-4 groups (**Figs. 4b** and **4c; Supp. Table 10**). In this context, these populations between groups showed that neutrophils and Treg/Th17 consistently increased in the treated groups, while macrophages grow more modestly (**Fig. 4d**). Thus, there is a strong association between neutrophils and Treg/Th17 (R = 0.92; *p* = 0.025), a positive but not significant relationship between macrophages and Treg/Th17 (R = 0.54; *p* = 0.35), and no significant correlation between neutrophils and macrophages (R = 0.22; *p* = 0.72).

**Figure 4.**
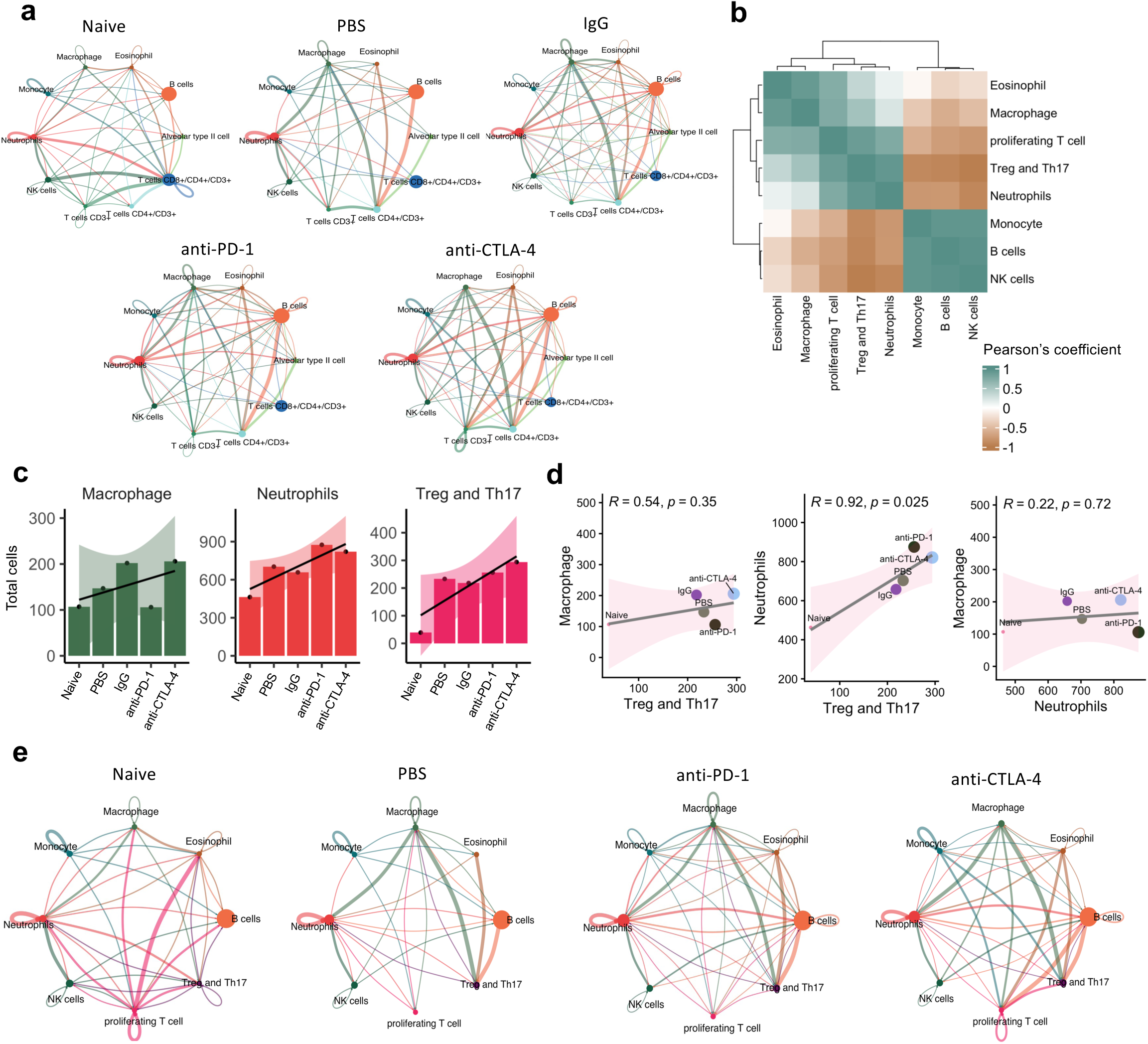
Checkpoint blockade strengthens Treg/Th17–neutrophil–macrophage communication in the infected lung. **a**) Global cell-cell communication networks derived from CellChat for the naïve, PBS, IgG, anti-PD-1, and anti-CTLA-4 groups. Each node represents a cell type identified in the lung, with size proportional to total interaction strength. Edges indicate predicted ligand–receptor interactions between cell types, with thickness proportional to communication intensity. **b**) Heatmap of Pearson correlation coefficients between the frequencies of Treg/Th17, neutrophils, macrophages, eosinophils, and other lineages. **c**) Barplots showing the total cell counts for macrophages, neutrophils, and Treg/Th17 cells among the treatment groups. The shadows represent the SE of the cell ratio among the treatment groups. **d**) Scatter plots show the relationship between Treg and Th17 abundances compared to macrophages and neutrophils, as well as, the correlation between neutrophils and macrophages, in the Naive, PBS, IgG, anti-PD-1, and anti-CTLA-4 groups. Each point represents a treatment group. The lines indicate the direction of correlation, with shaded areas corresponding to SE. R values (Pearson correlation coefficient) and p-values are shown. **e**) CellChat networks highlighting Treg/Th17 as dominant receivers of signals from macrophages and neutrophils in infected and treated groups, supporting a myeloid–Treg/Th17 axis that reorganizes the pulmonary microenvironment during infection and therapy. Each node represents a cell type identified in the lung, with size proportional to total interaction strength. Edges indicate predicted ligand–receptor interactions between cell types, with thickness proportional to communication intensity

In the PBS, anti-PD-1, and anti-CTLA-4 groups, compared to Naive, cell populations show that Treg/Th17 appears as a very evident receptor, consistent as a regulatory hub in *P. brasiliensis* infection (**Fig. 4e**). Additionally, interactions between these cells and macrophages and neutrophils were observed, indicating a natural inflammatory response to infection in which the neutrophil/macrophage (main emitters) and Treg/Th17 (receptor) axis is necessary to reorganize the pulmonary microenvironment during infection. Despite this response pattern during infection, cell communication between groups may diverge in L-R. Using the R NisheNet package^32^, we aimed to demonstrate in more detail the differences between the cell communication pattern between groups for the cell populations of neutrophils and macrophages (ligands) and Treg/Th17 (receptors). We performed a differential expression analysis between the Naive vs. PBS, PBS vs. anti-PD-1, and PBS vs. anti-CTLA-4 groups, where the results demonstrated divergences between communication among these populations in each group (**Fig. 5a**).

**Figure 5.**
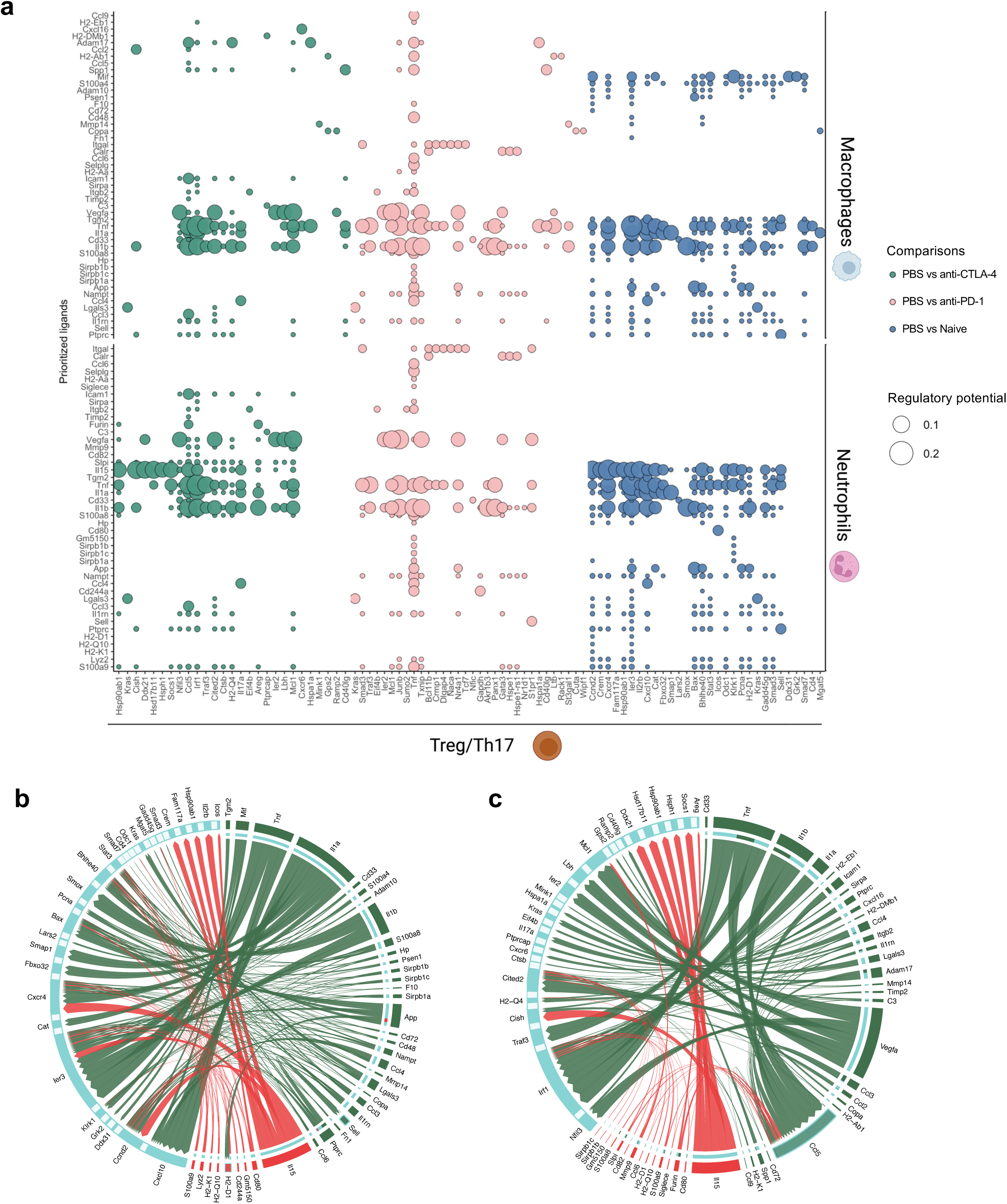
NicheNet reveals an *Il17a*-centered ligand–receptor axis linking myeloid cells to Treg/Th17 under CTLA-4 blockade. **a**) Bubble plot of NicheNet-derived ligand activities and regulatory potential scores for macrophage- and neutrophil-derived ligands targeting Treg/Th17 in Naïve vs PBS, PBS vs anti-PD-1, and PBS vs anti-CTLA-4 comparisons. The colors green, pink, and blue represent the comparison between the groups PBS vs. anti-CTLA-4, PBS vs. anti-PD-1, and Naive vs. PBS, respectively. The lines on the left represent predicted ligands for macrophages (above) and neutrophils (below), and the X-axis represents Treg/Th17 receptors. **b-c**) Circos plot for comparison of Naïve vs PBS (**b**) and PBS vs anti-CTLA-4 (**c**). The colors of nodes and edges in green and red indicate macrophage and neutrophil cell populations, respectively. In addition, the light blue nodes represent Treg/Th17 receptors.

Corroborating the results of the relative effects (**Fig. 3g**), *Il17a* and *Cxcr6* proved to be important molecules for Treg/Th17 in communication with macrophages and neutrophils in treatment with anti-CTLA-4. Among the emitters, macrophages and neutrophils similarly presented *Slpi*, *Ccl4*, *Il1a*, *Il1b*, *Tnf*, and *Il15* as ligands associated with *Il17a*, supporting a pro-inflammatory scenario ordered by treatment with anti-CTLA-4. Additionally, the *Cxcr6* target in Treg/Th17 showed interactions restricted to *Tnf* and *Cxcl16* exclusively in macrophages, suggesting the permanence of these cells in the pulmonary microenvironment. Furthermore, it is important to note that the neutrophil *Il15* ligand had *Il17a* from Treg/Th17 as an important target, suggesting maintenance and reorganization of these cells in the pulmonary microenvironment (**Fig. 5a**).

In summary, the data indicate that while infection (PBS) sustains a network focused on the mobilization and maintenance of neutrophils and macrophages, CTLA-4 blockade shifts communication toward *Il17a* (**Fig. 5b**-**5c** and **Supp. Table 11**). These findings define a specific deviation of anti-CTLA-4 compared to infection alone, with an increase in signals linked to *Il17a* and directed interaction between Treg/Th17 and neutrophil and macrophage cells in the pulmonary microenvironment, also suggesting an increase in the proinflammatory context, enhancing antifungal control via more effector neutrophils (**Fig. 6**).

**Figure 6.**
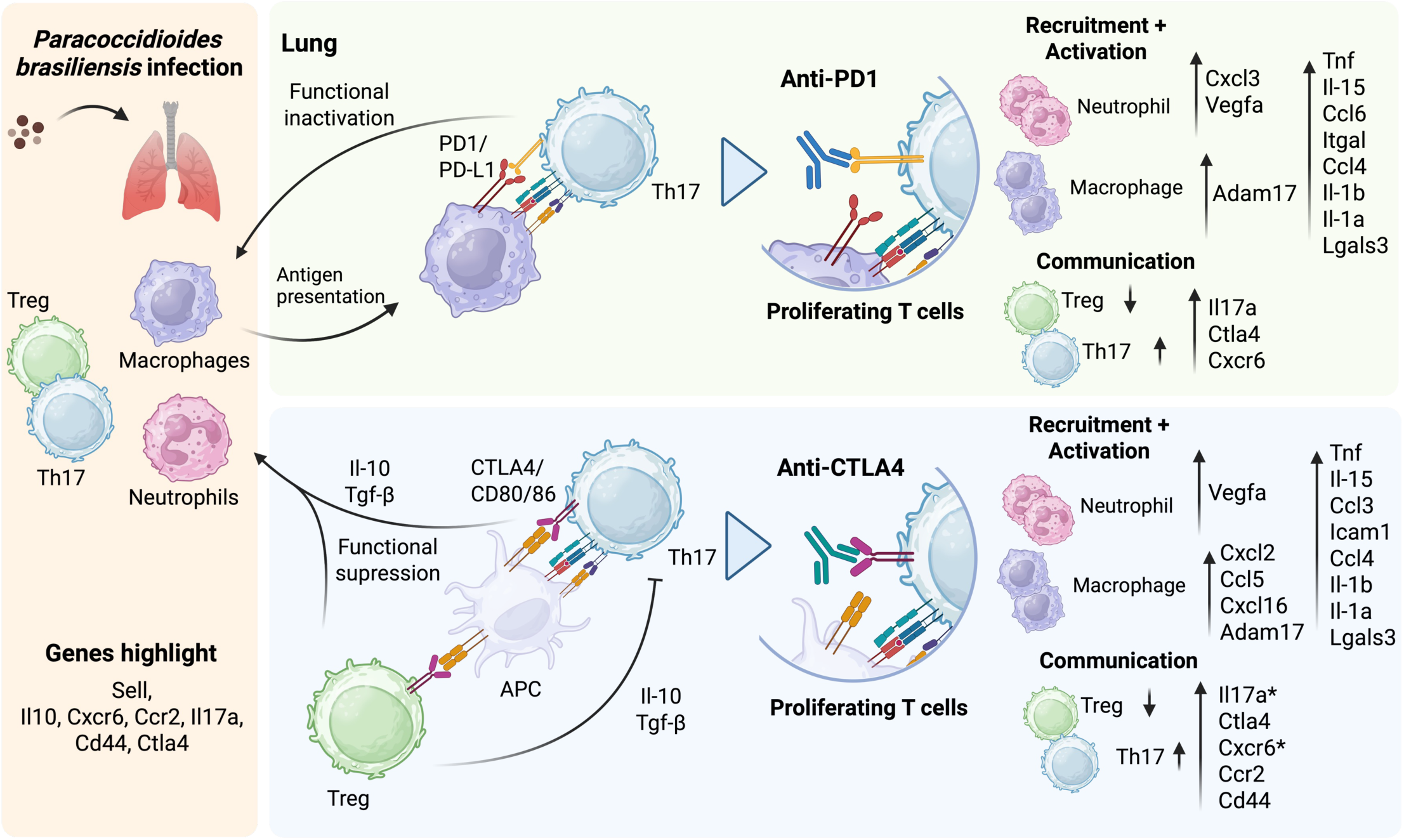
Proposed model of PD-1 and CTLA-4 blockade reshaping the Th17–Treg–myeloid circuit in pulmonary PCM. Schematic summary of *P. brasiliensis* lung infection showing baseline interactions between Treg, Th17, macrophages, and neutrophils and the distinct effects of PD-1 and CTLA-4 inhibition. In the PD-1 axis, blockade restores T cell activation and proliferation, enhances Th17 responses, and promotes recruitment/activation of neutrophils and macrophages via genes such as *Tnf*, *Il1b*, *Il15*, *Cxcl3*, *Cxcl5*, and *Vegfa*. In the CTLA-4 axis, blockade lifts suppressive signaling from Treg/APC interactions, amplifies Th17 differentiation and *Il17a* expression, and reinforces communication with macrophages and neutrophils through *Slpi*, *Ccl4*, *Il1a*, *Il1b*, *Tnf*, *Il15*, *Cxcr6*, and *Cxcl16*. The model highlights how checkpoint inhibition increases pro-inflammatory yet regulated effector circuits that support antifungal control while improving fungal elimination.

## DISCUSSION

Our findings reinforce the view that immune checkpoint pathways play a central role in defining the course and severity of *P. brasiliensis* infection, revealing that blocking PD-1 and CTLA-4 blockade can reorganize the pulmonary immune landscape in favor of fungal control. Previous studies have demonstrated that the balance between Th17 and Treg responses critically influences susceptibility and tissue damage in fungal infections, with IL-17-driven pathways promoting neutrophil recruitment and antifungal activity, while IL-10-producing Tregs facilitate pathogen dissemination and chronic disease^15,16,27,33–37^. In this context, our data extend previous observations by demonstrating that checkpoint inhibition not only decreases regulatory constraints but also enhances Th17-associated inflammatory mechanisms, a pattern that aligns with established IL-17 and neutrophil signaling mechanism^14,38^.

It is important to note that recent advances in anti-CTLA-4 and anti-PD-1 treatments highlight how modulation of inhibitory pathways can reprogram anti-tumor immunity^24,39–41^. These approaches, which selectively deplete suppressive Treg populations while enhancing effector responses, have demonstrated therapeutic efficacy with reduced toxicity in preclinical tumor models, underscoring the ability of checkpoint inhibition to recalibrate immune networks in vivo^39^. Parallel evidence from immunoregulatory research emphasizes the fundamental balance between Th17 and Treg cells in immune response effectiveness^37,42,43^; IL-17-induced pathways reinforce effective pathogen clearance, while excessive regulatory activity is associated with chronic disease states in multiple infection models^16,35,44,45^. These observations provide a stronger basis that aligns with our findings, reinforcing the idea that alleviating inhibitory activity can shift the pulmonary immune landscape toward a more effective response against *P. brasiliensis*^15,16,35,42,44,46^.

Building on this framework, the regulation of the Th17 and Treg cell axis is determinant for fungal dissemination has been well described^33^, and studies in both PCM patients and experimental models show that IL-17 together with IFN-γ and TNF is essential for effective immunity, whereas the expansion of IL-10–producing Treg correlates with more severe and disseminated disease^20,47,48^. Our previous work using this infection model demonstrated that blockade of CTLA-4 or PD-1 reduces fungal burden in multiple organs and markedly limits tissue damage, suggesting that inhibiting inhibitory signaling enhances the host’s capacity to control *P. brasiliensis* infection^26^. The scRNA-seq data presented here reinforce this interpretation: treatment with either antibody shifted the pulmonary immune environment toward a Th17-skewed profile accompanied by reduction in *Il10*-expressing Treg and a corresponding reprogramming of neutrophil and macrophage responses^15,26,38,43,46,48–50^. This integrative response aligns with the improved fungal clearance observed previously^26^ and supports a mechanistic model in which increased IL-17-mediated activation and recruitment of neutrophils and macrophages directly contribute to fungal containment^14,20,48,51^.

Checkpoint inhibition further amplified the effector profile of Th17 cells, which displayed reduced exhaustion and increased expression of *Il17a*, *Ccr2,* and *Cxcr6*. This shift is consistent with the well-established role of IL-17A in directing neutrophil recruitment to sites of infection^38,49^, and aligns with the marked infiltration of neutrophils and macrophages we observed in the treated lung. The transcriptomic signatures indicate Th17-mediated communication that helps maintain effector function^50^. Within this environment, interactions between Treg/Th17 and myeloid populations were characterized by the concurrent expression of mediators such as *Il17*, *Slpi*, *Ccl4*, *Il1a*, *Il1b*, *Tnf* and *Il15*, as highlighted by our NicheNet analysis^32^. These signals point to an immune microenvironment that becomes progressively more permissive to myeloid activation once checkpoint pathways are blocked^14,52–57^.

Consistent with this, anti-CTLA-4 and anti-PD-1 treatment induced robust TNF-α expression and a transcriptional program linking Th17 cells, neutrophils, and macrophages^26,58^. Clinical and experimental data show that Th1-derived IFN-γ and TNF-α act synergistically to drive fungicidal activity in macrophages, while therapeutic blockade of TNF-α exacerbates fungal disease severity^20,26,47^. Our results reflect these observations and highlight a strengthened inflammatory axis involving Th17, Th1, and myeloid populations. In addition, checkpoint blockade also attenuated Treg-associated^35^ suppressive pathways, including reductions in *Il10*, *Foxp3*, *Ctla4*, and *Pdcd1*, thereby releasing regulatory constraints known to limit antifungal immunity^59–61^. Thus, checkpoint inhibition reorganizes the pulmonary immune landscape into a state more conducive to *P. brasiliensis* clearance, consistent with previous reports of reduced fungal disease burden and tissue pathology after anti-PD-1 or anti-CTLA-4 treatment^26^.

Several lines of evidence help contextualize our observations. Experimental models have shown that disruption of IL-17–associated pathways, including IL-6, IL-23, and IL-17RA signaling, impairs fungal control and alters granuloma organization^12^, while IDO1 deficiency leads to exaggerated Th17 activity and insufficient Treg-mediated counter-regulation^33^. Across fungal infections, IL-1α, IL-1β, and TNF serve as central mediators of early inflammatory responses but can also contribute to tissue injury^62^. Moreover, differences in virulence among *P. brasiliensis* strains have been associated with distinct neutrophil activation pathways, ranging from balanced TNF-α/IL-10 responses to highly proinflammatory programs driven by TNF-α, GM-CSF, IL-1 and reactive oxygen species^63–66^. Additionally, our ligand-receptor analysis identified *Il15* within the neutrophil interaction network, and its documented role in antifungal immunity, through natural killer (NK) and T-cell regulation, further supports the relevance of our findings^62^.

In light of this framework, our single-cell analysis reveals how PD-1 and CTLA-4 blockade restructures the Th17–Treg–myeloid axis during *P. brasiliensis* infection. Checkpoint inhibition enhanced Th17–macrophage communication, reduced Treg suppressive signatures, and promoted neutrophil and macrophage activation, generating a transcriptional landscape consistent with effective fungal containment. Although PD-1 and CTLA-4 blockade produced overlapping outcomes, each therapy elicited a distinct transcriptional signature, underscoring their differential roles in immune regulation. Together, these findings corroborate previous observations based on cytometry and demonstrate that immune checkpoint inhibition improves coordinated cellular communication between Th17, Treg, neutrophils, and macrophages. Importantly, this immune activity occurs without evidence of exacerbated inflammatory responses, supporting a balanced immunomodulatory effect rather than uncontrolled immune activation during antifungal immunity in PCM.

## METHODS

### Study design

C57BL/6 mice were intratracheally infected with 1×10⁶ viable *P. brasiliensis* Pb18 yeast cells in 50 µL PBS, following established experimental paracoccidiodomycosis protocols^26^. After three weeks of infection, a time point at which adaptive immunity and regulatory checkpoint pathways are firmly established, animals received intraperitoneal treatment with immune checkpoint inhibitors. Mice were injected three times per week with 200 µg of either anti-CTLA-4 (clone UC10F10-11) or anti-PD-1 (clone RMP1-14), corresponding to approximately 10 µg/kg, or received 200 µg of isotype-matched IgG or PBS. Treatments were maintained for five consecutive weeks. At the end of this period, corresponding to the eight-week post-infection, animals were euthanized, pulmonary leukocytes were isolated, and a transcriptomic single-cell RNA sequencing was performed for evaluation of immune responses.

### Ethical Statement

All animal procedures were conducted in strict accordance with institutional and national ethical guidelines for animal experimentation. Protocols were approved by the Institutional Animal Care and Use Committee (IACUC) under the corresponding project license (protocol number CEUA 180/11). Mice were anesthetized with intraperitoneal injections of xylazine (30 mg/kg) and ketamine (180 mg/kg) and euthanized at 8 weeks post-infection or earlier if signs of pain or distress (e.g., loss of appetite, lethargy, arching of the spine, or piloerection) were observed.

### Lung Cell Preparation

Lungs were harvested after mice euthanasia and enzymatically digested with 1 mg/mL of collagenase type IV (Sigma Aldrich, CAT#C4-22-1G, USA) for 30 min under agitation for 30 min at 37 °C. Cell suspensions were separated on Percoll (Cytiva, CAT#17089101) gradient (33%), centrifuged at 900 × g for 10 min at 4 °C, filtered through a 70 µm cell strainer (Corning, CAT#431751, USA), and treated with BD ACK Lysing Buffer (BD Biosciences, CAT#555899, USA) to remove red blood cells. Cells were counted and stained using the BD™ Mouse Immune Single-Cell Multiplexing Kit (BD Biosciences, CAT#633793, USA) targeting CD45 for 20 min at room temperature, labeling 1 × 10⁶ cells per mouse across the uninfected, Pb+PBS, Pb+IgG, Pb+anti-CTLA4, and Pb+anti-PD1 groups (total of 4 sample tags). Stained cells were washed, labeled for viability using Calcein-AM (2 mM, Invitrogen, CAT#C1430, USA) and Draq7 (0.3 mM, BD Biosciences, CAT#564904, USA) for 5 min at 37 °C, and counted with Trypan Blue (Invitrogen, CAT#15250061, USA) to assess viability.

### Single-Cell Capture, Library Construction, and Sequencing

Freshly isolated lung cells were processed immediately using the BD Rhapsody™ scRNA-seq platform (BD Biosciences). Five pooled samples (uninfected, Pb+PBS, Pb+IgG, Pb+anti-CTLA4, and Pb+anti-PD1; total ∼55,000 cells) were loaded per BD Rhapsody™ cartridge kit (BD Biosciences, CAT#633733, USA), using four cartridges in total (20 samples). Across all groups, 70,474 cells were recovered (∼14,094 cells/group).

Single-cell capture was performed in microwells (20 min at room temperature) followed by cell lysis and poly-A mRNA hybridization to barcoded magnetic beads, as per the manufacturer’s instructions. After retrieval, reverse transcription introduced unique molecular identifiers (UMIs) to each cDNA molecule. Whole-transcriptome amplification (WTA) and sample-tag libraries were generated using the BD Rhapsody™ Single-Cell WTA Amplification Kit (BD Biosciences, CAT#633801, USA) and purified with Agencourt® AMPure® XP beads (Beckman Coulter, CAT#A63880, USA). Library quality and quantity were verified using the Qubit dsDNA HS Assay Kit (Invitrogen) and Agilent 4200 TapeStation. Pooled libraries (4 nM initial dilution) were denatured and sequenced on an Illumina NovaSeq 6000 system (Illumina) using a NovaSeq 6000 S1 Flow Cell (Illumina, CAT# 20028318, USA) (2 × 75 bp, no PhiX) at ∼20,000 reads/cell and 600 reads/sample tag.

### Single cell RNA Sequence analysis

The sequencing data (CBCL format) were processed with bcl2fastq^67^ to generate FASTQ files. We used the Salmon-Alevin^68–71^ software to map and quantify the reads against the reference transcriptome and genome for *Mus musculus*^70^. The analyses were conducted using the R programming language and RStudio (R version 4.5.0 (2025-04-11))^71^. The objects were constructed with the Seurat^69^ package from the raw count matrices, with quality filtering per cell based on the number of genes detected, number of UMIs, and mitochondrial fraction (<20%). To avoid clustering biases caused by tags, after merging the samples into their respective groups, the sample tags were removed from the analyses. In addition, according to the study design, the samples were normalized and corrected for batch effects with NormalizeData, FindVariableFeatures, and ScaleData, followed by dimensionality reduction by principal component analysis (PCA). The neighborhood structure was estimated with FindNeighbors, and clusters were obtained via FindClusters, with resolution chosen by topological stability and biological separation using the R clustree package^72^.

### Cell cluster annotation by enrichment and differential expression gene analysis

Using the Seurat^31^ and presto packages, we annotated the profile of each cluster based on differential markers (FindAllMarkers) and functional enrichment analysis with the R clusterprofiler^73^ package using the comparecluster function. The biological processes (BPs) used for annotation were filtered based on an adjusted p-value (False Discovery Rate [FDR]) < 0.05.

### Wilcoxon test for cell cluster

To test for differences in abundance per cluster between groups, we calculated the proportion of cells per sample and used 95% confidence interval (CI) values to estimate significant increases in cell populations. In addition, we compared the absolute value of cell counts per group using the Wilcoxon test. P-values were adjusted by Benjamini–Hochberg (BH-FDR), and we reported test statistics, adjusted p-values, and effects. Statistical significance was considered with adjusted p-values < 0.05.

### Trajectory analysis and Relative Effect with MANOVA

Cell pseudotime inference was performed using the R Slingshot^74^ package. This approach allows the reconstruction of cell differentiation pathways based on gene expression data in reduced dimensionality space^75,76^. Initially, cells were grouped using unsupervised clustering, and UMAP coordinates were used as input for the trajectory inference algorithm. Pseudotime analysis was performed using subcluster 2 (initial trajectory cluster) for cluster 1 (final trajectory cluster). This configuration was used for the different groups (Naive, PBS, IgG, anti-PD1, and anti-CTLA4).

Furthermore, using differentially expressed genes (DEGs), we evaluated the relative effects of specific genes in the comparisons Naive vs. PBS, PBS vs. IgG, PBS vs. anti-PD1, and PBS vs. anti-CTLA4. The relative effects were estimated using multivariate analysis of variance (MANOVA), employing a bootstrap approach with 1,000 resamples to increase the reliability of the results. This strategy allowed us to consider the inherent variability in the data and calculate confidence intervals for the observed effects, providing a statistically robust estimate of gene associations. All analyses were conducted in the R environment, using the npmv^77^ and reshape2^78^ packages, such as described^79^.

### Cell Chat and Pearson’s correlation

To characterize cellular interaction patterns in the pulmonary microenvironment, we performed cellular communication analysis using the CellChat^80^ package in the R environment. The analyses used the CellChatDB^81^ Mouse database as a reference. Initially, the complete dataset was analyzed considering all populations together with all groups normalized together. Next, we conducted a targeted analysis focused on specific populations, including CD4⁺ T cells subpopulations. In addition, correlations were calculated using the *cor* function in R base, employing Pearson’s method to assess the linear relationship between variables.

### NicheNet analysis

To identify upstream ligands potentially responsible for transcriptional changes in recipient cell populations, a ligand-target inference analysis was performed using the NicheNet R package^32^. Macrophages and neutrophils were defined as sender cells and Treg/Th17 as receivers. Differential gene expression analysis was performed on the recipient cell population, comparing the groups Naive vs PBS, PBS vs anti-PD1, and PBS vs anti-CTLA4 within each population. Genes with *p* ≤ 0.05 and |log2 fold change| ≥ 0.25 were selected as the set of genes of interest. This set of genes was crossed with the NicheNet ligand-target matrix to ensure compatibility with prior knowledge. Background-expressed genes were defined as genes detected in the recipient population and present in the ligand-target matrix. Ligand activities were predicted using the predict_ligand_activities function, and ligands were ranked based on the area under the precision-recall curve (AUPR). The 30 ligands with the highest predicted activity were selected for further analysis. Weighted ligand-target interactions were inferred using get_weighted_ligand_target_links, retaining the top 100 target genes per ligand. Ligand-target links were filtered using a weight cutoff of 0.33 and used for network visualization and further interpretation.

## Code availability

Code will be made publicly available upon publication. All R packages used in this study are described in the manuscript.

## Data availability

All data supporting the findings of this study are available within the paper and its Supplementary Information. Raw scRNA-seq data will be deposited in the Sequence Read Archive (SRA) upon acceptance of the manuscript.

## Contributions

D.L.M.F., I.A.F.B., and S.M.M. wrote the manuscript; D.L.M.F., I.A.F.B., S.M.M., P.S., L.M.G., E.S., J.M.Z., C.A.T., K.T.K., M.O.R.S., C.N.C.L., B.P.E., and L.S.B.M. provided scientific insights; D.L.M.F. and P.S. performed data and bioinformatics analyses; D.L.M.F., I.A.F.B., D.M.F., F.V.L., V.L.G.C., and S.M.M. revised and edited the manuscript; N.W.P., B.M.B., V.L.K., M.C.A., B.V.S., D.M.F., F.V.L.., and S.M.M. performed the experiments and analyzed the data ; V.L.G.C. conceived and designed the project, and provided funds to carry out the project. S.M.M., D.L.M.F. and I.A.F.B. contributed equally and are considered co-first authors.

## Acknowledgments

We acknowledge the São Paulo Research Foundation (FAPESP grants: 2024/08802-3 and 2024/08360-0 S.M.M.; 2025/03433-2 to D.L.M.F.; 2025/04633-5 to C.A.T.; 2024/06780-2 to C.N.C.L.; 2022/06074-5 to E.S.; 2025/10345-2 to B.P.E.; 2023/10839-0 to L.S.B.M.; 2021/06881-5 to D.M.F.; 2022/10275-6 to L.M.G.; 2024/14290-5 to B.C.O.; 2019/12691-4 to M.C.A.; 2022/08700-0 and 2024/06233-1 to J.M.Z; 2023/15407-0 to F.V.L; and 2020/08460-4 to V.L.G.C.), and the Coordination for the Improvement of Higher Education Personnel (CAPES) Financial Code 88887.992612/2024-00 (grant to I.A.F.B) for financial support. We acknowledge the National Council for Scientific and Technological Development (CNPq) Brazil grant 303384/2024-7 to S.M.M.

## Supplementary Figures legends

**Supplementary Figure S1. Relative treatment effects on cell cluster abundance.** Forest plot of odds ratios and 95% confidence intervals summarizing the significant increase in overall cell count for each cluster and between the Naïve, PBS, IgG, anti-PD-1, and anti-CTLA-4 groups as a whole. The purple and red dots indicate significant decreases and increases. In contrast, the gray dots represent non-significant changes.

**Supplementary Figure S2. Marker gene expression supporting lineage annotation of lung clusters.** UMAP feature plots for representative lineage and activation markers (e.g., *Cd3e*, *Cd4*, *Cd8a*, *Il17a*, *Ifng*, *Ly6g*, *Cd19*, *Trem2*, *S100a9*, *Ear2*, *Hbb-bt*, *Mpeg1*, *Sftpb*), illustrating their spatial distribution across the 11 clusters.

**Supplementary Figure S3. Reclustering and characterization of total T cells in infected and treated lungs. a**) UMAP of reclustered CD3^+^ T cells showing five transcriptionally distinct T subpopulations. **b**) Feature plots for T cell markers used to define CD8^+^, mixed CD4^+^/CD8^+^, and CD4^+^ helper phenotypes. **c**) Violin plots of key markers across T subclusters, indicating activated cytotoxic, helper, and proliferative states. **d**) Bar plots of T subcluster frequencies per group, demonstrating expansion of CD4^+^ subsets in anti-PD-1 and anti-CTLA-4 conditions. The shadows represent the standard error (SE).

**Supplementary Figure S4. Transcriptional features of Treg/Th17 and proliferative CD4^+^ T subclusters. a**) UMAP feature plots showing expression of Treg and Th17 markers (*Foxp3*, *Il10*, *Il17a*, *Rora*, *Ccr2*, *Cxcr6*, *Cd44*) in CD4^+^ T subclusters. **b**) Feature plots for proliferation-associated genes (*Mki67*, *Top2a*, *Pcna*, *Stmn1*) highlighting the proliferative cluster 2. **c**) Additional markers associated with activation and migration (*Ccr7*, *Cxcr3*, *Gzmb*, *Klrk1*, *Itgax*) across CD4^+^ states. **d**) Violin plots of Il17a and key genes per group within the clusters, showing increased Il17a expression in anti-PD-1 and anti-CTLA-4 compared to PBS. **e**) Feature plots illustrating treatment-specific modulation of selected genes, supporting differentiation from proliferative CD4^+^ T cells toward Treg/Th17 phenotypes under checkpoint blockade.

## REFERENCES

1. Nucci, M., Colombo, A. L. & Queiroz-Telles, F. Paracoccidioidomycosis. Curr. Fungal Infect. Rep. 3, 15–20 (2009).

2. Ekeng, B. E. et al. Pulmonary and Extrapulmonary Manifestations of Fungal Infections Misdiagnosed as Tuberculosis: The Need for Prompt Diagnosis and Management. Journal of Fungi 8, 460 (2022).

3. Ashraf, N. et al. Re-drawing the Maps for Endemic Mycoses. Mycopathologia 185, 843–865 (2020).

4. Turissini, D. A., Gomez, O. M., Teixeira, M. M., McEwen, J. G. & Matute, D. R. Species boundaries in the human pathogen Paracoccidioides. Fungal Genetics and Biology 106, 9–25 (2017).

5. Colombo, A. L., Tobón, A., Restrepo, A., Queiroz-Telles, F. & Nucci, M. Epidemiology of endemic systemic fungal infections in Latin America. Med. Mycol. 49, 785–798 (2011).

6. Tirado-Sánchez, A., González, G. M. & Bonifaz, A. Endemic mycoses: epidemiology and diagnostic strategies. Expert Rev. Anti. Infect. Ther. 18, 1105–1117 (2020).

7. Calich, V. L., Vaz, C. A. & Burger, E. Immunity to Paracoccidioides brasiliensis infection. Res. Immunol. 149, 407–417; discussion 499–500 (1998).

8. Queiroz-Telles, F. V. de, Peçanha Pietrobom, P. M., Rosa Júnior, M., Baptista, R. de M. & Peçanha, P. M. New Insights on Pulmonary Paracoccidioidomycosis. Semin. Respir. Crit. Care Med. 41, 53–68 (2020).

9. Blair, J. E. et al. Characteristics of patients with mild to moderate primary pulmonary coccidioidomycosis. Emerg. Infect. Dis. 20, 983–990 (2014).

10. Spinello, I. M., Munoz, A. & Johnson, R. H. Pulmonary coccidioidomycosis. Semin. Respir. Crit. Care Med. 29, 166–173 (2008).

11. Ye, P. et al. Requirement of interleukin 17 receptor signaling for lung CXC chemokine and granulocyte colony-stimulating factor expression, neutrophil recruitment, and host defense. J. Exp. Med. 194, 519–527 (2001).

12. Tristão, F. S. M. et al. Th17-Inducing Cytokines IL-6 and IL-23 Are Crucial for Granuloma Formation during Experimental Paracoccidioidomycosis. Front. Immunol. 8, (2017).

13. Mota, N. G. S. et al. Correlation between cell-mediated immunity and clinical forms of paracoccidioidomycosis. Trans. R. Soc. Trop. Med. Hyg. 79, 765–772 (1985).

14. Meloni-Bruneri, L. H. et al. Neutrophil oxidative metabolism and killing of *P. brasiliensis* after air pouch infection of susceptible and resistant mice. J. Leukoc. Biol. 59, 526–533 (1996).

15. Loures, F. V., Pina, A., Felonato, M. & Calich, V. L. G. TLR2 Is a Negative Regulator of Th17 Cells and Tissue Pathology in a Pulmonary Model of Fungal Infection. The Journal of Immunology 183, 1279–1290 (2009).

16. de Araújo, E. F., Preite, N. W., Veldhoen, M., Loures, F. V. & Calich, V. L. G. Pulmonary paracoccidioidomycosis in AhR deficient hosts is severe and associated with defective Treg and Th22 responses. Sci. Rep. 10, 11312 (2020).

17. de Araújo, E. F. et al. AhR Ligands Modulate the Differentiation of Innate Lymphoid Cells and T Helper Cell Subsets That Control the Severity of a Pulmonary Fungal Infection. Front. Immunol. 12, (2021).

18. Chiarella, A. P. et al. The relative importance of CD4+ and CD8+T cells in immunity to pulmonary paracoccidioidomycosis. Microbes Infect. 9, 1078–1088 (2007).

19. Cano, L. E. et al. Depletion of CD8 + T Cells In Vivo Impairs Host Defense of Mice Resistant and Susceptible to Pulmonary Paracoccidioidomycosis. Infect. Immun. 68, 352–359 (2000).

20. Bernardino, S. et al. TNF-α and CD8+ T Cells Mediate the Beneficial Effects of Nitric Oxide Synthase-2 Deficiency in Pulmonary Paracoccidioidomycosis. PLoS Negl. Trop. Dis. 7, e2325 (2013).

21. Cardoso, R. M. N. et al. The relation between FoxP3 + regulatory T cells and fungal density in oral paracoccidioidomycosis: a preliminary study. Mycoses 57, 771–774 (2014).

22. PD-1 Blockade with Nivolumab in Relapsed or Refractory Hodgkin’s Lymphoma New England Journal of Medicine. Preprint at https://www.nejm.org/doi/full/10.1056/NEJMoa1411087.

23. Gao, Y. et al. Acetylation-dependent regulation of PD-L1 nuclear translocation dictates the efficacy of anti-PD-1 immunotherapy. Nat. Cell Biol. 22, 1064–1075 (2020).

24. Seidel, J. A., Otsuka, A. & Kabashima, K. Anti-PD-1 and Anti-CTLA-4 Therapies in Cancer: Mechanisms of Action, Efficacy, and Limitations. Front. Oncol. 8, 86 (2018).

25. Campanelli, A. P. et al. Fas-Fas ligand (CD95-CD95L) and cytotoxic T lymphocyte antigen-4 engagement mediate T cell unresponsiveness in patients with paracoccidioidomycosis. J. Infect. Dis. 187, 1496–1505 (2003).

26. Preite, N. W. et al. Blocking the CTLA-4 and PD-1 pathways during pulmonary paracoccidioidomycosis improves immunity, reduces disease severity, and increases the survival of infected mice. Front. Immunol. 15, (2024).

27. Galdino, N. A. L. et al. Depletion of regulatory T cells in ongoing paracoccidioidomycosis rescues protective Th1/Th17 immunity and prevents fatal disease outcome. Sci. Rep. 8, 16544 (2018).

28. Angelidis, I. et al. An atlas of the aging lung mapped by single cell transcriptomics and deep tissue proteomics. Nat. Commun. 10, 963 (2019).

29. Ansell, S. M. et al. PD-1 Blockade with Nivolumab in Relapsed or Refractory Hodgkin’s Lymphoma. New England Journal of Medicine 372, 311–319 (2015).

30. Na, Z. et al. Structural basis for blocking PD-1-mediated immune suppression by therapeutic antibody pembrolizumab. Cell Res. 27, 147–150 (2017).

31. Hao, Y. et al. Dictionary learning for integrative, multimodal and scalable single-cell analysis. Nat. Biotechnol. 42, 293–304 (2024).

32. Browaeys, R., Saelens, W. & Saeys, Y. NicheNet: modeling intercellular communication by linking ligands to target genes. Nat. Methods 17, 159–162 (2020).

33. de Araújo, E. F. et al. The IDO–AhR Axis Controls Th17/Treg Immunity in a Pulmonary Model of Fungal Infection. Front. Immunol. Volume 8–2017, (2017).

34. Mills, K. H. G. IL-17 and IL-17-producing cells in protection versus pathology. Nat. Rev. Immunol. 23, 38–54 (2023).

35. Calich, V. L. G., Mamoni, R. L. & Loures, F. V. Regulatory T cells in paracoccidioidomycosis. Virulence 10, 810–821 (2019).

36. Ferreira, M. C., De Oliveira, R. T. D., Da Silva, R. M., Blotta, M. H. S. L. & Mamoni, R. L. Involvement of Regulatory T Cells in the Immunosuppression Characteristic of Patients with Paracoccidioidomycosis. Infect. Immun. 78, 4392–4401 (2010).

37. Preite, N. W., Kaminski, V. de L., Borges, B. M., Calich, V. L. G. & Loures, F. V. Myeloid-derived suppressor cells are associated with impaired Th1 and Th17 responses and severe pulmonary paracoccidioidomycosis which is reversed by anti-Gr1 therapy. Front. Immunol. 14, (2023).

38. Gaffen, S. L. Structure and signalling in the IL-17 receptor family. Nat. Rev. Immunol. 9, 556–567 (2009).

39. Cao, W. et al. A next-generation anti-CTLA-4 probody mitigates toxicity and enhances anti-tumor immunity in mice. Nat. Commun. 16, 9029 (2025).

40. Babamohamadi, M. et al. Anti-CTLA-4 nanobody as a promising approach in cancer immunotherapy. Cell Death Dis. 15, 17 (2024).

41. Ward, F. J., Kennedy, P. T., Al-Fatyan, F., Dahal, L. N. & Abu-Eid, R. CTLA-4—two pathways to anti-tumour immunity? Immunotherapy Advances 5, ltaf008 (2024).

42. Araújo, E. F. de et al. Tolerogenic Plasmacytoid Dendritic Cells Control Paracoccidioides brasiliensis Infection by Inducting Regulatory T Cells in an IDO-Dependent Manner. PLoS Pathog. 12, e1006115 (2016).

43. Loures, F. V., AraÃojo, E. F., Feriotti, C., Bazan, S. B. & Calich, V. L. G. TLR-4 cooperates with Dectin-1 and mannose receptor to expand Th17 and Tc17 cells induced by Paracoccidioides brasiliensis stimulated dendritic cells. Front. Microbiol. 6, (2015).

44. Bazan, S. B., et al. Loss- and Gain-of-Function Approaches Indicate a Dual Role Exerted by Regulatory T Cells in Pulmonary Paracoccidioidomycosis. PLoS Negl. Trop. Dis. 9, e0004189 (2015).

45. Whibley, N. & Gaffen, S. L. Brothers in Arms: Th17 and Treg Responses in Candida albicans Immunity. PLoS Pathog. 10, e1004456 (2014).

46. Loures, F. V. et al. Dectin-1 Induces M1 Macrophages and Prominent Expansion of CD8+IL-17+ Cells in Pulmonary Paracoccidioidomycosis. J. Infect. Dis. 210, 762–773 (2014).

47. Cano, L. E. et al. Protective Role of Gamma Interferon in Experimental Pulmonary Paracoccidioidomycosis. Infect. Immun. 66, 800–806 (1998).

48. Costa, T. A., et al. In Pulmonary Paracoccidioidomycosis IL-10 Deficiency Leads to Increased Immunity and Regressive Infection without Enhancing Tissue Pathology. PLoS Negl. Trop. Dis. 7, e2512 (2013).

49. Lin, I.-C. et al. Involvement of IL-17 A/IL-17 Receptor A with Neutrophil Recruitment and the Severity of Coronary Arteritis in Kawasaki Disease. J. Clin. Immunol. 44, 77 (2024).

50. Xing, J., Man, C., Liu, Y., Zhang, Z. & Peng, H. Factors impacting the benefits and pathogenicity of Th17 cells in the tumor microenvironment. Front. Immunol. 14, (2023).

51. Kashino, S. S. et al. Effect of macrophage blockade on the resistance of inbred mice toParacoccidioides brasiliensis infection. Mycopathologia 130, 131–140 (1995).

52. Bruns, S. et al. Production of Extracellular Traps against Aspergillus fumigatus In Vitro and in Infected Lung Tissue Is Dependent on Invading Neutrophils and Influenced by Hydrophobin RodA. PLoS Pathog. 6, e1000873 (2010).

53. Loures, F. V. et al. Recognition of Aspergillus fumigatus Hyphae by Human Plasmacytoid Dendritic Cells Is Mediated by Dectin-2 and Results in Formation of Extracellular Traps. PLoS Pathog. 11, e1004643 (2015).

54. Zhang, F. et al. Neutrophil diversity and function in health and disease. Signal Transduct. Target. Ther. 9, 343 (2024).

55. Urban, C. F. & Nett, J. E. Neutrophil extracellular traps in fungal infection. Semin. Cell Dev. Biol. 89, 47–57 (2019).

56. Rogan, M. P. et al. Antimicrobial proteins and polypeptides in pulmonary innate defence. Respir. Res. 7, 29 (2006).

57. Brown, G. D. et al. Hidden Killers: Human Fungal Infections. Sci. Transl. Med. 4, 165rv13–165rv13 (2012).

58. Tsiodras, S., Samonis, G., Boumpas, D. T. & Kontoyiannis, D. P. Fungal Infections Complicating Tumor Necrosis Factor α Blockade Therapy. Mayo Clin. Proc. 83, 181–194 (2008).

59. Diep, A. L. & Hoyer, K. K. Host Response to Coccidioides Infection: Fungal Immunity. Front. Cell. Infect. Microbiol. 10, 581101 (2020).

60. Neworal, E. Immunocytochemical localization of cytokines and inducible nitric oxide synthase (iNOS) in oral mucosa and lymph nodes of patients with paracoccidioidomycosis. Cytokine 21, 234–241 (2003).

61. Pagliari, C. et al. Paracoccidioidomycosis: Cells expressing IL17 and Foxp3 in cutaneous and mucosal lesions. Microb. Pathog. 50, 263–267 (2011).

62. Caffrey-Carr, A. K. et al. Interleukin 1α Is Critical for Resistance against Highly Virulent Aspergillus fumigatus Isolates. Infect. Immun. 85, e00661–17 (2017).

63. Bonfim, C. V., Mamoni, R. L. & Blotta, M. H. S. L. TLR-2, TLR-4 and dectin-1 expression in human monocytes and neutrophils stimulated by Paracoccidioides brasiliensis. Med. Mycol. 47, 722–733 (2009).

64. Rodrigues, D. R., Dias-Melicio, L. A., Calvi, S. A., Peraçoli, M. T. S. & Soares, A. M. V. C. *Paracoccidioides brasiliensis* killing by IFN-γ, TNF-α and GM-CSF activated human neutrophils: role for oxygen metabolites. Med. Mycol. 45, 27–33 (2007).

65. Tavian, E. G. et al. Interleukin-15 increases Paracoccidioides brasiliensis killing by human neutrophils. Cytokine 41, 48–53 (2008).

66. Nishikaku, A. S. et al. Nitric oxide participation in granulomatous response induced by Paracoccidioides brasiliensis infection in mice. Med. Microbiol. Immunol. 198, 123–135 (2009).

67. Luecken, M. D. & Theis, F. J. Current best practices in single-cell RNA-seq analysis: a tutorial. Mol. Syst. Biol. 15, (2019).

68. Srivastava, A., Malik, L., Sarkar, H. & Patro, R. A Bayesian framework for inter-cellular information sharing improves dscRNA-seq quantification. Bioinformatics 36, i292–i299 (2020).

69. Srivastava, A., Malik, L., Smith, T., Sudbery, I. & Patro, R. Alevin efficiently estimates accurate gene abundances from dscRNA-seq data. Genome Biol. 20, 65 (2019).

70. Church, D. M. et al. Modernizing Reference Genome Assemblies. PLoS Biol. 9, e1001091 (2011).

71. Wickham, H. et al. Welcome to the Tidyverse. J. Open Source Softw. 4, 1686 (2019).

72. Zappia, L. & Oshlack, A. Clustering trees: a visualization for evaluating clusterings at multiple resolutions. Gigascience 7, giy083 (2018).

73. Yu, G., Wang, L.-G., Han, Y. & He, Q.-Y. clusterProfiler: an R Package for Comparing Biological Themes Among Gene Clusters. OMICS 16, 284–287 (2012).

74. Street, K. et al. Slingshot: cell lineage and pseudotime inference for single-cell transcriptomics. BMC Genomics 19, 477 (2018).

75. Rostom, R., Svensson, V., Teichmann, S. A. & Kar, G. Computational approaches for interpreting scRNA-seq data. FEBS Lett. 591, 2213–2225 (2017).

76. Lyu, M. et al. Single-Cell Sequencing Reveals Functional Alterations in Tuberculosis. Advanced Science 11, 2305592 (2024).

77. Burchett, W. W., Ellis, A. R., Harrar, S. W. & Bathke, A. C. Nonparametric Inference for Multivariate Data: The R Package npmv. J. Stat. Softw. 76, 1–18 (2017).

78. Wickham, H. Reshaping Data with the reshape Package. J. Stat. Softw. 21, 1–20 (2007).

79. Fonseca, D. L. M. et al. Dysregulated autoantibodies targeting AGTR1 are associated with the accumulation of COVID-19 symptoms. NPJ Syst. Biol. Appl. 11, 7 (2025).

80. Jin, S., Plikus, M. V & Nie, Q. CellChat for systematic analysis of cell–cell communication from single-cell transcriptomics. Nat. Protoc. 20, 180–219 (2025).

81. Jin, S. et al. Inference and analysis of cell-cell communication using CellChat. Nat. Commun. 12, 1088 (2021).

