## Supplementary figures and images for "Single-cell RNA sequencing of CTLA-4 and PD-1 blockade in pulmonary paracoccidioidomycosis highlights a protective transcriptional program mediated by activated Th17 cells, neutrophils and macrophages"

### Supplementary Figures

**Supp. figure 1**

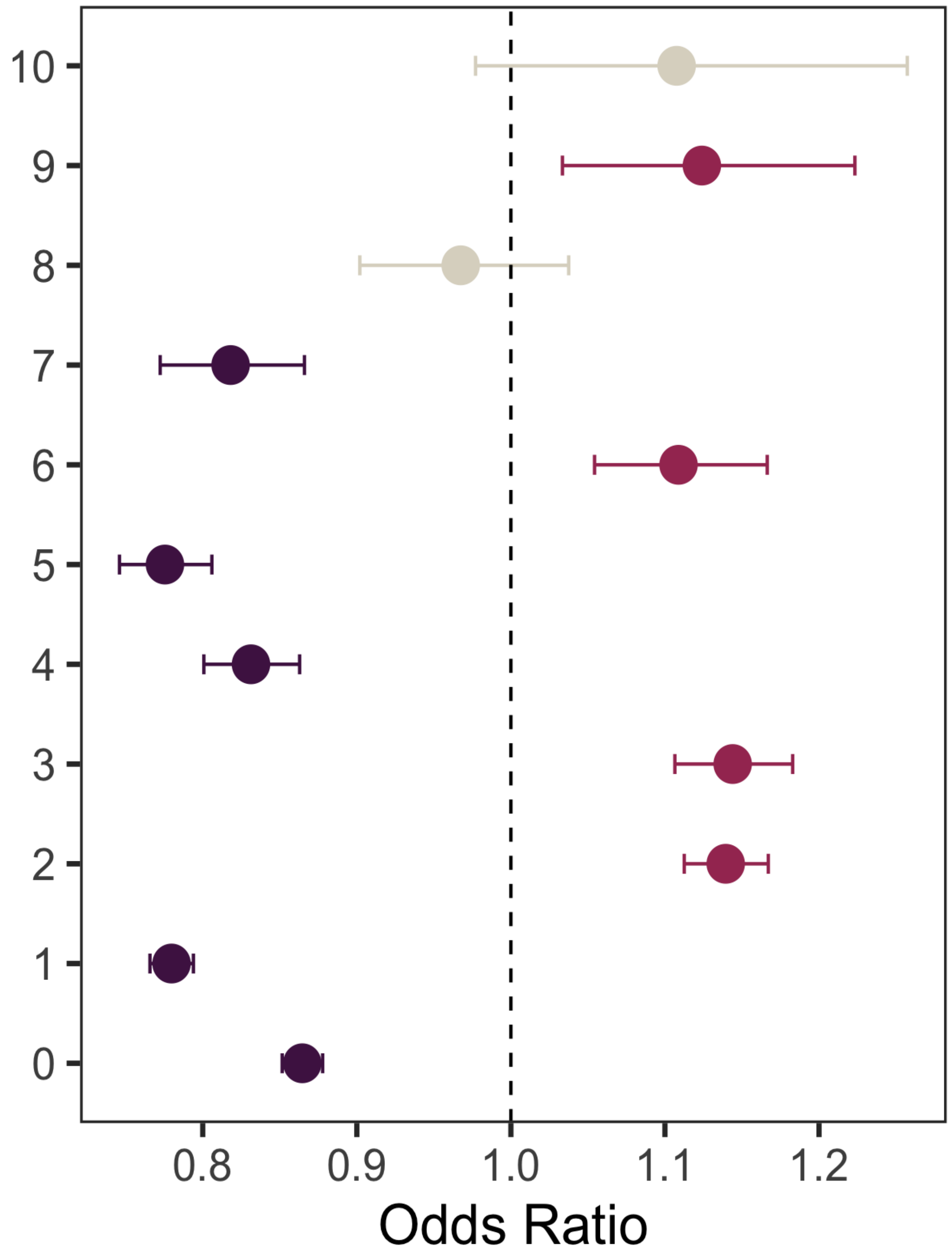

**Supp. figure 2**

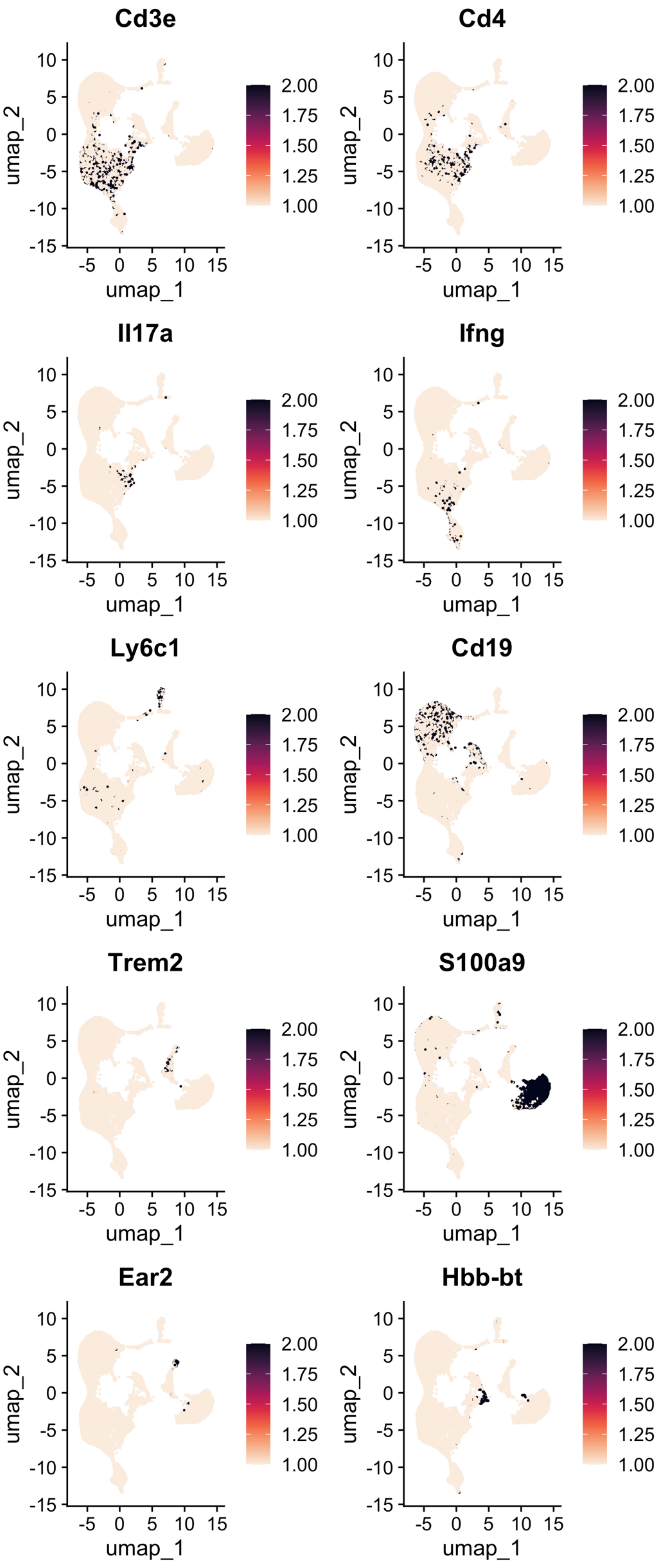

Supp. figure 3

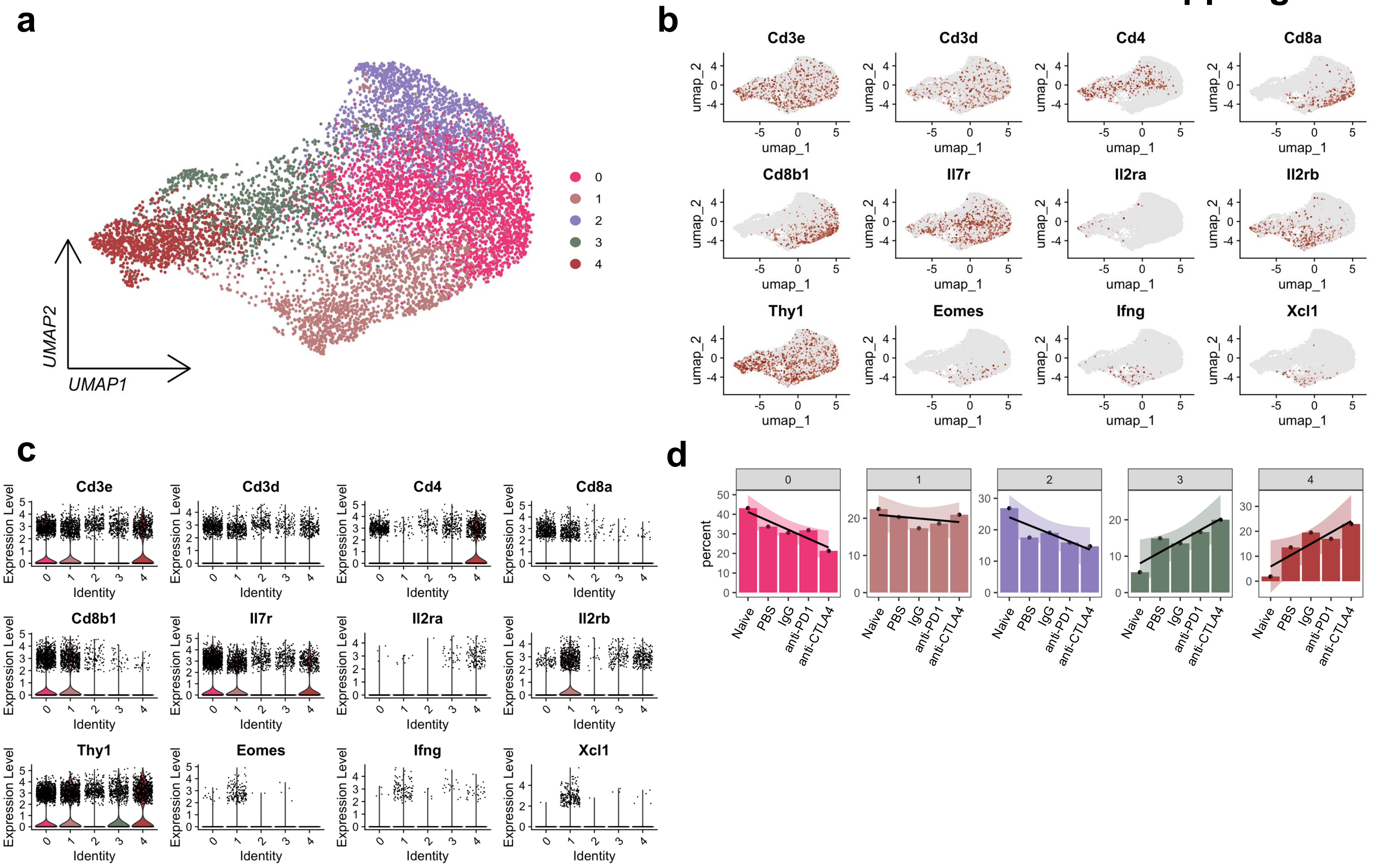

#### Supp. figure 4

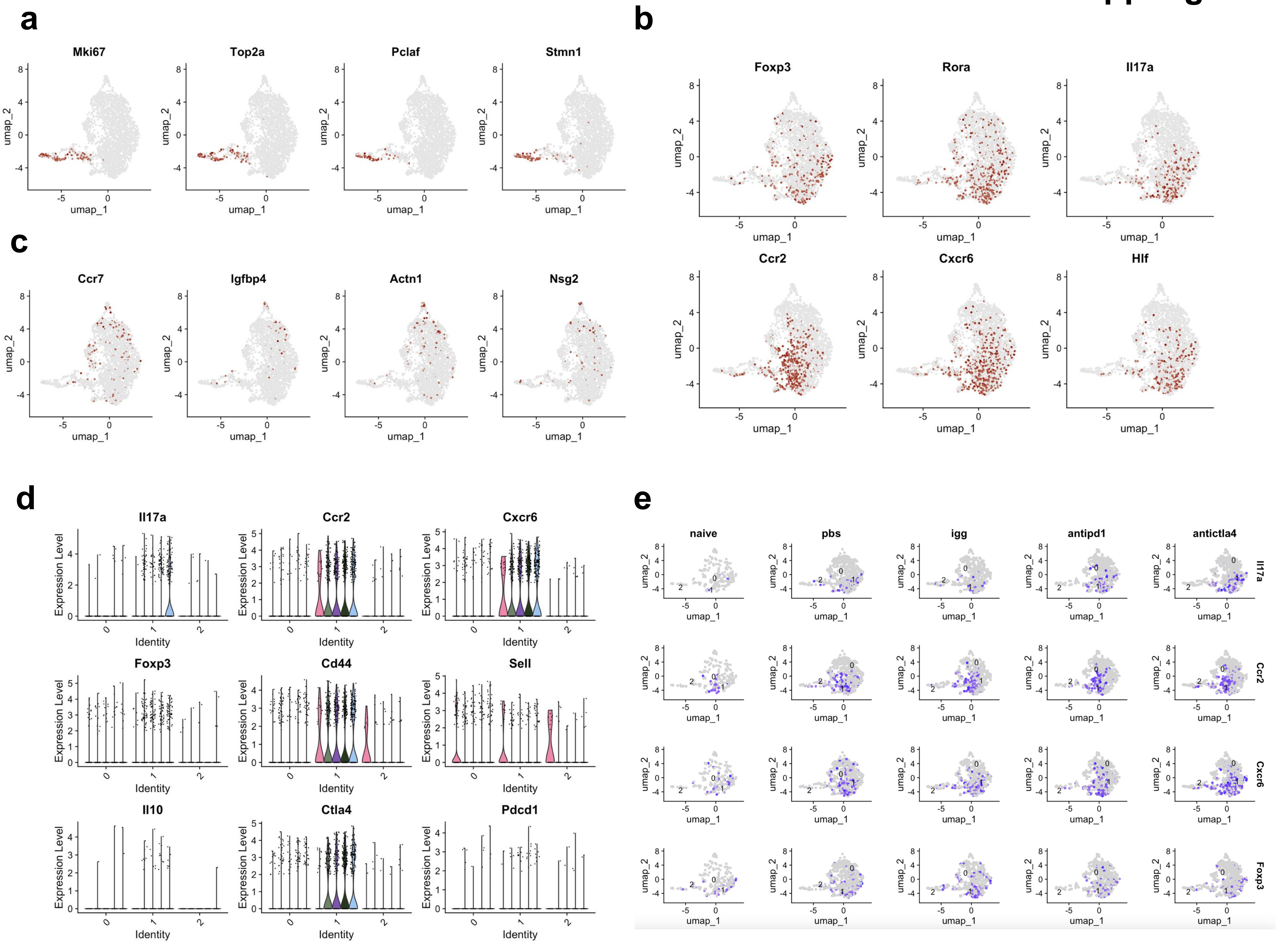
